# Collapse of local circuit integrated information Φ during NREM sleep

**DOI:** 10.64898/2026.04.01.715799

**Authors:** Keiichi Onoda

## Abstract

Clarifying the mechanisms underlying the emergence of consciousness remains a fundamental challenge in modern neuroscience. Integrated Information Theory (IIT) provides a mathematical framework derived from the phenomenological properties of consciousness as its axioms. IIT proposes that consciousness is identical to a system’s intrinsic cause-effect information structure, quantified by integrated information Φ. While IIT predicts that the Φ of a neuronal system should decrease during the loss of consciousness, this hypothesis has remained untested at the neural circuit level. The present study provides empirical support for this IIT prediction. It was found that Φ within local circuits decreases during non-rapid eye movement (NREM) sleep compared to wakefulness and REM sleep, regardless of cortical laminar structure or brain regions, and even after controlling for firing rates. The reduction in Φ was particularly pronounced during OFF-periods, when neural activity is collectively suppressed. These results align with IIT’s framework, suggesting that the loss of consciousness corresponds to a collapse of integrated information within local circuits that cannot be explained solely by changes in individual system elements (such as overall firing rates). The findings provide empirical support for IIT as a mathematical theory aiming to explain conscious experiences.

## Introduction

The quest to identify the mechanisms underlying the emergence of consciousness remains one of the most challenging issues in neuroscience. While numerous neurobiological theories of consciousness have been proposed (Northoff and Lamme 2020), Integrated Information Theory (IIT) is distinguished by its unique phenomenology-to-physics approach (Tononi 2004). Central to this framework is the explanatory identity, which posits that a specific conscious experience is identical to the cause-effect information structure unfolded from a physical substrate. According to IIT, consciousness is not merely a process, but a fundamental property of physical systems that possess intrinsic, irreducible cause-effect power. For a system to be conscious, it must specify a cause-effect repertoire that is at least highly informative (selecting one specific state out of many possibilities) and highly integrated (causal power exists only as a unified whole). The quantity of consciousness is measured by integrated information (Φ). In IIT 4.0 (Albantakis et al. 2023), Φ quantifies an intrinsic causal power of the system, specifically measuring the degree to which the system’s ability to influence its own past and future cannot be reduced to the independent contributions of its parts. If a system’s causal power is lost upon partitioning, it possesses high Φ, signifying a high degree of conscious existence.

Empirical investigations of Φ using neural activity remain limited, especially at the micro-level. Hypotheses derived from IIT have primarily been tested by applying various proxy measures of Φ to macro-level activity (Baglivo et al. 2024; Kim et al. 2018; Luppi et al. 2019; Luppi et al. 2024; Massimini et al. 2005; Onoda and Akama 2023; Onoda and Akama 2024). However, these macro-level proxies might not distinguish between informational richness and causal information integration (Mediano et al. 2022). Theoretical developments in IIT suggest that integrated causal power is not necessarily maximized at the level of individual elements (i.e., the micro-scale, such as single neurons) but may emerge at an appropriate spatiotemporal grain (i.e., the macro-scale, such as groups of neurons) through coarse-graining and black-boxing (Hoel et al. 2013; Hoel et al. 2016; Marshall et al. 2018; Marshall et al. 2024). In fact, empirical findings corroborate this, demonstrating that Φ is maximized not at micro-timescales, but at specific macro-timescales (Leung and Tsuchiya 2022). Recently, based on this insight, formal Φ measures have been applied to coarse-grained macro-level brain activity, with studies consistently reporting a reduction in Φ during the loss of consciousness associated with anesthesia and sleep (Leung et al. 2021; Nemirovsky et al. 2023; Onoda et al. 2025). However, a critical methodological gap remains. The systems defined in IIT assume irreducible elements (Albantakis et al. 2023), a criterion that continuous and spatially mixed signals like local field potentials (LFPs) and blood oxygenation level-dependent (BOLD) activity used in these studies do not fully satisfy. Therefore, to rigorously align empirical data with IIT’s theoretical framework, it is necessary to construct relatively macro-level systems from neuronal spikes, which better approximate the genuine irreducible units of information processing, and to validate the changes in Φ within such systems.

Sleep serves as a natural experiment for testing the predictions of IIT. While rapid eye movement (REM) dreaming resembles wakeful experience, non-REM (NREM) sleep often involves a loss of consciousness (Tononi et al. 2024). Indeed, dreaming reports are less frequent in NREM compared to REM sleep (Siclari et al. 2017). The hallmark of NREM sleep is the slow wave, reflecting a cortical bistability where neuronal populations switch between ON-periods and silent OFF-periods (Steriade et al. 1993; Steriade et al. 2001). From an IIT perspective, this bistability underlies the loss of consciousness (Tononi and Cirelli 2020). The emergence of OFF-periods effectively disrupts causal interactions between neurons (Kiyooka et al. 2026; Senzai et al. 2019), decomposing the neural substrate and signaling a loss of integration. Sustained inactivity during OFF-periods collapses the repertoire of available states, representing a loss of information (González et al. 2021; Massimini et al. 2010; Tononi et al. 2024). These findings suggest that cortical Φ decreases during NREM sleep, particularly within OFF-periods, and recovers during REM sleep. To investigate these dynamics, the present study constructed coarse-grained systems from cortical neural spikes and evaluated the associated changes in Φ.

## Materials and Methods

The datasets used in this study were downloaded from Buzsáki Lab webshare (https://buzsakilab.nyumc.org/datasets/SenzaiY/) (Petersen et al. 2020; Senzai et al. 2019) and Collaborative Research in Computational Neuroscience (https://crcns.org/data-sets/fcx/fcx-1) (Watson et al. 2016). In this study, these are referred to as the V1 and FC datasets, respectively. These two datasets were selected because they provide neural population activities across wakefulness, NREM sleep, and REM sleep.

### Datasets

**V1**: Nineteen male mice were used. They were implanted with 64-channel single-shank silicon probes targeting the primary visual cortex (V1), oriented perpendicular to the cortical surface to align with apical dendrites. Extracellular signals were recorded at 20 kHz while the mice behaved freely in their home cages. A single recording session was conducted for each mouse, with a duration of 244 ± 100 min. Single units were sorted and identified using Kilosort. The number of isolated units recorded per session averaged 77.7 ± 21.6 (range: 38–119). Brain states (wakefulness, NREM, and REM) were categorized via principal component analysis of LFP spectrograms, narrowband theta power ratios, and EMG signals extracted from LFP correlations. Five physiological landmarks were established to align cortical depth profiles across animals.

**FC**: Eleven male Long-Evans rats were used. Each rat was implanted with one or two 64-site silicon probes targeting frontal cortical areas, including the medial prefrontal, anterior cingulate, secondary motor, and orbitofrontal cortices. Single-unit activity and LFPs were recorded at 20 kHz while rats were in their home cages. Recording sessions lasted an average of 294 ± 137 min. Spikes were clustered using the KlustaKwik algorithm. Across 27 recording sessions, this study identified an average of 41.5 ± 24.0 stable units per session (range: 10–113). Brain states (wakefulness, NREM, REM, and microarousals) were automatically categorized using principal component analysis of LFP spectrograms, narrowband theta power ratios, and EMG measures. Locomotor activity of the subjects was also recorded in the FC dataset.

### Experimental Design

An overview of the data analysis pipeline is illustrated in Fig. 1. Both datasets include neural population activity and state information (wakefulness, NREM, and REM sleep) determined by local field potentials (Fig. 1A). Three conditions (WAKE, NREM, and REM) were categorized in accordance with the labels provided in the original dataset. The target systems were constructed as follows. For the V1 dataset, the available cortical depth information was leveraged (Fig. 1B & C) to define a system consisting of layers as elements (Fig. 1D1). Additionally, systems for both datasets were constructed by randomly assigning units into elements (Fig. 1D2). The activities of all units within each element were aggregated into a single cluster (Fig. 1E). To identify the system states, the presence or absence of spikes was determined within each cluster for every specified time window with duration τ (Fig. 1F). Specifically, multiple spike trains were integrated to define the elements of the system, and extended τ values were employed to construct a coarse-grained system. Based on the time series of these states, a transition probability matrix (TPM) was estimated (Fig. 1G), from which the Φ values for each state were calculated (Fig. 1H). Finally, these Φ values were mapped back onto the state time series (Fig. 1I), and the mean Φ for each condition was calculated for each session. Details of each step are described below.

**Figure 1.**
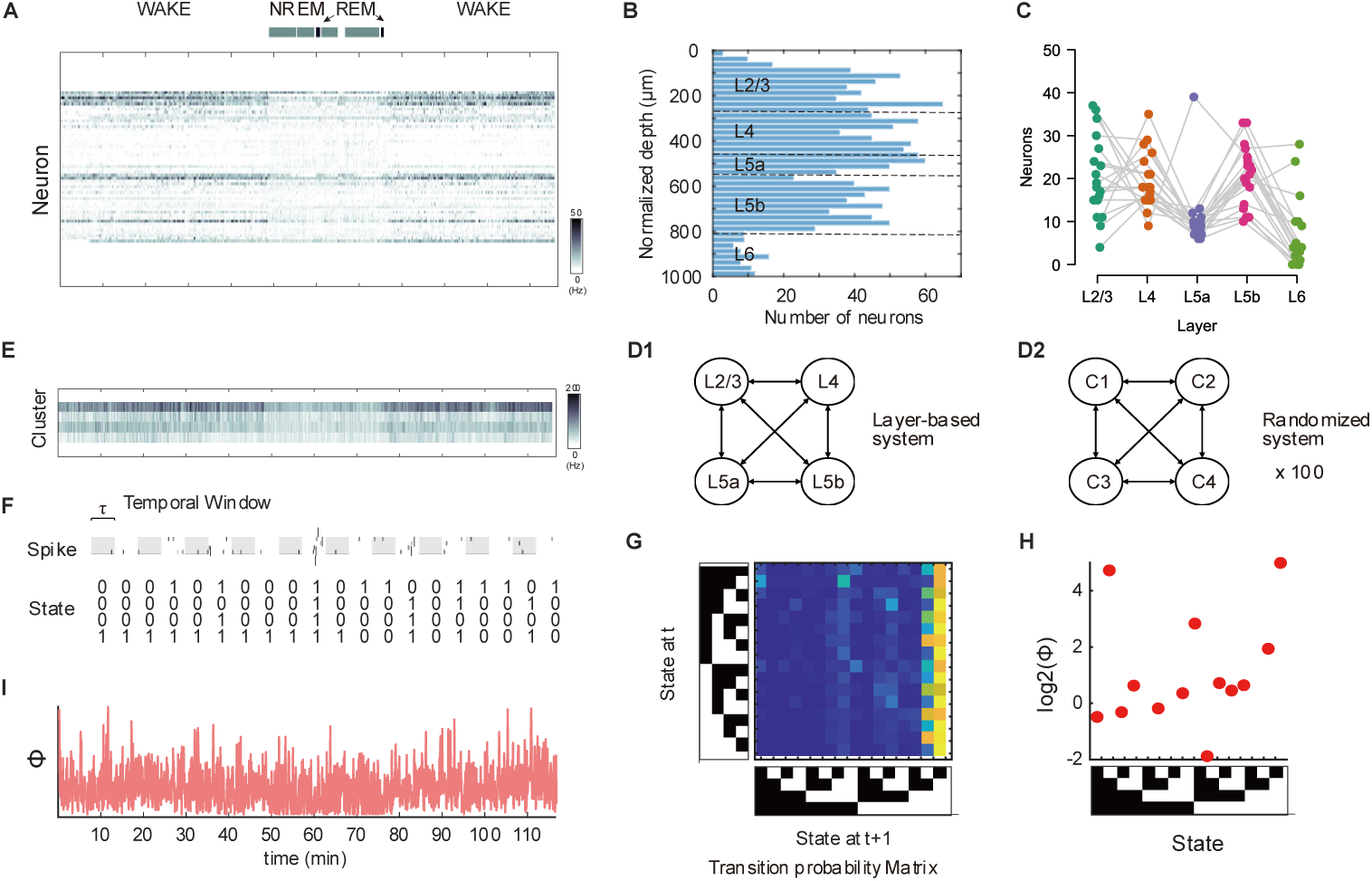
Analysis workflow. (A) Raster plot of a single session including WAKE, NREM, and REM epochs. (B) Frequency distribution of neurons by cortical depth. (C) Frequency distribution of neuron counts per cortical layer for each session. (D1&D2) Construction of layer-based and randomized systems. (E) Aggregation of neural activity into elements. (F) Binary signals were generated over discrete time using a window τ. The state sequence was obtained from the binary signals of the four selected elements. (G&H) The transition probability matrix (TPM) was computed from the state sequence, and Φ for each state was computed from the TPM. (I) The Φ value for each state was assigned to the corresponding state in the sequence, and the values were averaged in each condition.

### Unit selection

The calculation of Φ is computationally intensive, with memory and processing time requirements increasing exponentially as the number of system elements grows. Consequently, previous studies have typically evaluated Φ in systems composed of 4 to 5 elements. In this study, the number of system elements was fixed at four.

For the V1 dataset, standardized cortical depth and layer identity are provided for each unit. As shown in Fig. 1B and C, units were roughly uniformly distributed across the depth range of 100 to 800 μm, with a sufficient number of units recorded in layers L2/3, L4, L5a, and L5b. Many sessions contained no units in L6; L2/3, L4, L5a, and L5b were selected as the four elements of the system (Fig. 1D1). All units within a given layer were aggregated into a single spike train, resulting in four layer-specific spike trains (Fig. 1E) that were used for the IIT analyses.

Furthermore, to investigate whether Φ depends on the laminar structure, a system independent of cortical layers was constructed. Specifically, units were randomly partitioned into four clusters, and spikes from units within each cluster were aggregated to form four spike trains. The system composed of these clusters as elements was also subjected to IIT analysis (Fig. 1D2). This randomized assignment approach was applied to both the V1 and FC datasets, with results averaged over 100 independent iterations.

### State time series

For each element of the system, a value of 1 or 0 was assigned depending on whether at least one neural spike occurred within a given time window (τ) (Fig. 1E&F). A value of 1 was assigned if any neuron within the layer or cluster fired, otherwise, 0 was assigned. The value of τ ranged from 10 to 100 ms, in steps of 10 ms in the V1 dataset. In the FC dataset, the value of τ ranged from 20 to 300 ms, in steps of 20 ms to alleviate the computational burden. The state at a given time point was represented as a combination of the element values (1 or 0) within the system, resulting in 2^4^ = 16 possible patterns. The states were indexed according to the little-endian convention. For example, state [1, 0, 0, 0] was assigned index 1 and [0, 1, 0, 0] was assigned index 2.

### Transition probability matrix

In a system where each element assumes a binary value, a TPM is essential for quantifying the level of integrated information. The TPM represents the probability of transitioning from one state to another (including self-transitions) at a subsequent time point. The frequencies of all observed transitions were used to construct a 16 × 16 square matrix (Fig. 1G). To normalize the matrix, each row was divided by the total number of transitions originating from the corresponding state (i.e., the number of times the state appeared in the time series and transitioned to any state). The TPM was calculated using the states observed during each condition (excluding periods of movement in the FC dataset).

### Integrated information Φ

In this study, Φ was computed according to IIT version 4 (Albantakis et al. 2023) using the PyPhi toolbox (Mayner et al. 2018). First, a TPM provides the fundamental probabilities of how the system transitions between states. Based on the TPM, the algorithm identifies the core system, which is the specific set of elements that is most strongly integrated and cannot be reduced into independent parts. Next, the algorithm analyzes this core system to find “distinctions” and “relations”. Distinctions represent the causal components within the system, while relations describe how these components overlap and interact. Each distinction and relation is assigned an integrated information value (φ), which measures its irreducibility based on the original TPM probabilities. Finally, the total structure integrated information, or “big Φ”, is calculated simply by summing the φ values of all distinctions and relations. Φ was calculated for all system states (Fig. 1H) and assigned to each time point according to the system’s state at that time (Fig. 1I). The average Φ for each condition was computed within each session by averaging Φ across time points, which corresponds to a weighted average across states based on their occurrence frequency.

## Controls for TPM estimation

The mean total durations of each condition (WAKE, NREM, REM) were 75.3 ± 52.2, 118.7 ± 58.8, and 12.1 ± 7.7 min in the V1 dataset, 33.5 ± 23.8, 76.6 ± 57.0, and 14.6 ± 13.6 min for the FC dataset, indicating a markedly shorter duration for REM sleep. Shorter durations of a given state may preclude the estimation of robust TPMs. To address this limited sampling problem, analyses were conducted using both condition-specific TPMs and a common TPM derived from the baseline (WAKE) condition. In the former approach, each time point in a state time series was assigned the state-wise Φ value derived from the TPM corresponding to the condition to which that time point belonged; in contrast, in the latter approach, the state-wise Φ value derived from the baseline TPM was assigned across all conditions. In addition, an analysis was performed where TPMs were constructed after equalizing the data length across conditions to match that of the condition with the shortest duration, from which Φ was recalculated for comparison. For this length equalization, continuous temporal segments centered around the median time point of each condition were extracted.

Furthermore, to verify whether condition-dependent differences in Φ were reliant on the structure of state transitions, TPMs were calculated from state time series whose temporal sequences were randomly shuffled, and the resulting Φ values were compared.

### Controls for firing rates

The firing rate per element was defined as the mean across the 4 elements of the total spike count within each element divided by the state duration and the number of constituent neurons. To test whether state-dependent changes in Φ were driven by firing rates, correlations between firing rates per element and Φ were evaluated across pooled data, individual conditions, and within-subject differences relative to WAKE. To eliminate firing rate as a confounding factor, mean firing rates across conditions within each subject were equalized by randomly downsampling spike events in higher-rate conditions to match the lowest-rate condition. Φ was then recalculated from the rate-equalized time series.

### Statistical analysis

In the V1 dataset, the Φ values were primarily analyzed using a repeated-measures analysis of variance (ANOVA) for each τ, with condition specified as the independent variable. Similarly, ANOVAs were performed on the maximum Φ, the optimal τ, and the firing rate per element for each condition. When conducting ANOVAs across multiple τ values, the p-values for the main effects were corrected for the number of τ levels using the Bonferroni method. Post hoc comparisons between the conditions were performed using Bonferroni corrected *p*-values. All *p*-values reported in the Results section are adjusted.

Statistical analyses for the FC dataset were performed using linear mixed-effects (LME) models to account for the repeated-measures structure and nested experimental design across subjects and recording sessions. For each dependent variable (e.g., Φ, firing rates), an LME model was constructed with condition and brain region as fixed effects. To account for subject-level baseline differences and inter-session variability, nested random intercepts were included for subjects and recording sessions nested within subjects (1|Subject and 1|Subject:Session). Main effects of the fixed factors were evaluated using Type III ANOVA. When a main effect of condition was identified, post-hoc comparisons between conditions were conducted using linear contrast tests. The *p*-values for post-hoc comparisons were adjusted for multiple testing using the Bonferroni correction.

## Results

### Layer-based systems in V1 dataset

Fig. 2 shows the results of Φ for each analysis setting in layer-based systems in the V1 dataset. The top panels present the results based on condition-specific TPMs, whereas the bottom panels present those based on a common TPM (WAKE). Fig. 2A illustrates Φ as a function of τ for each condition. Across conditions, Φ was highest around τ = 50 ms, with lower values observed in NREM and REM compared to WAKE. Comparing the τ yielding the maximum Φ across conditions (Fig. 2B; *F(2,36)* = 317.2, *η_p_^2^* > .95, *p* < .001) revealed that τ was significantly shorter in NREM than in WAKE and REM (*ps* < .002). Furthermore, a comparison of the maximum Φ across conditions (Fig. 2C; *F(2,36)* = 58.3, *η_p_^2^* = .76, *p* < .001) showed that Φ in NREM was significantly lower than in WAKE (*p* = .040). Next, to eliminate potential bias arising from differences in state sequence length across conditions, the analysis using length-equalized states was conducted and showed consistent results (Fig. 2D). Furthermore, to confirm whether the reduction of Φ in NREM depends on the underlying structure of state transitions, TPMs from temporally-shuffled state sequences were constructed and used for calculating Φ. Consequently, Φ was severely diminished to near-zero levels across all conditions (Fig. 2E).

**Figure 2.**
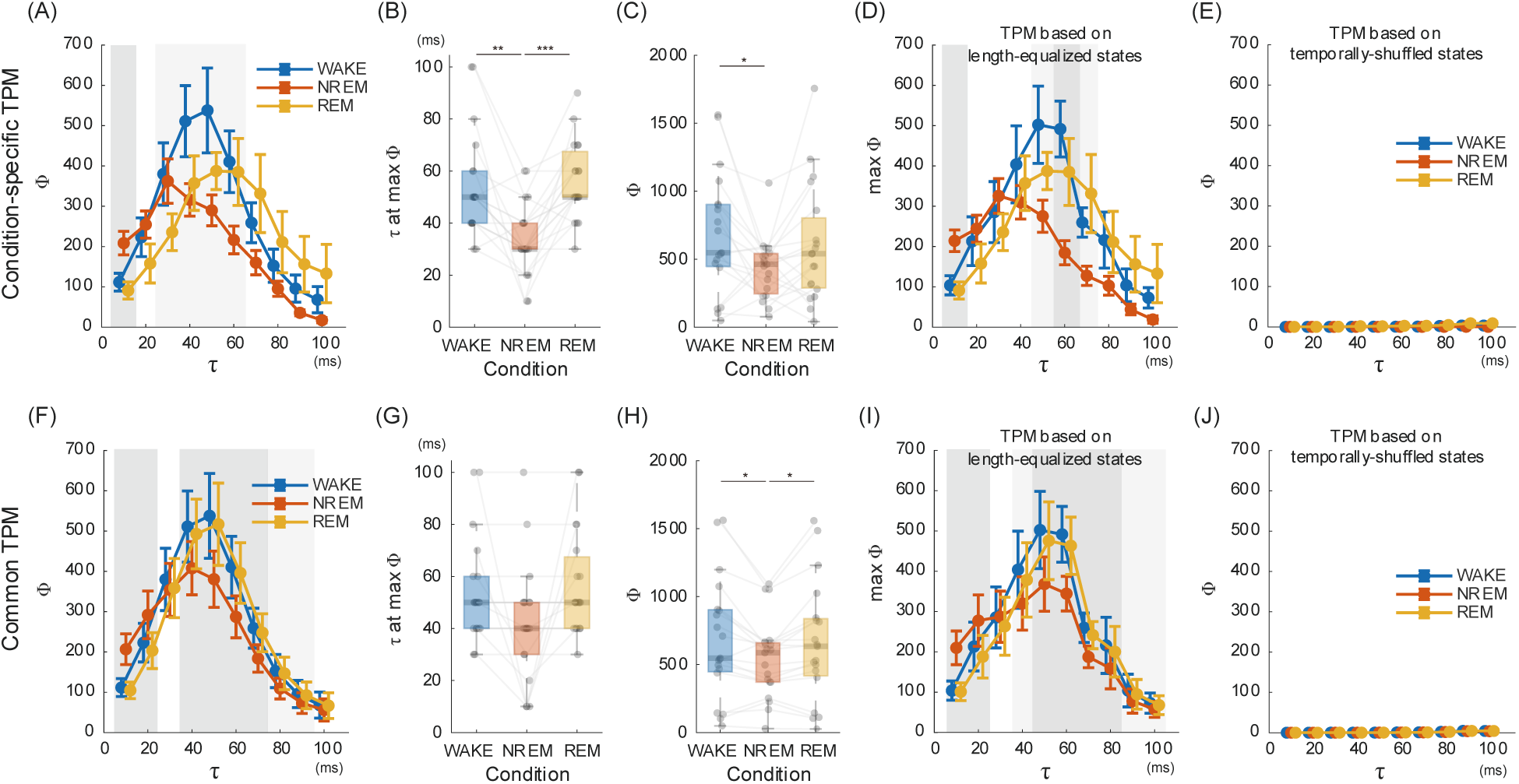
Φ in layer-based systems for the V1 dataset. Top panels show results where condition-specific transition probability matrices (TPMs) were used to calculate Φ for each condition. Bottom panels show results where Φ was calculated using the common TPM (WAKE). (A, F) Mean Φ as a function of τ across conditions. Error bars represent the standard error of the mean. Gray shaded areas indicate τ values with a significant main effect of condition, as determined by independent ANOVAs (dark gray: Bonferroni-corrected *p* < .05, light gray: uncorrected *p* < .05). (B, G) The τ yielding the maximum Φ for each condition. (C, H) Maximum Φ for each condition. (D, I) Mean Φ as a function of τ across conditions for systems with TPMs based on length-equalized states. (E, J) Mean Φ as a function of τ across conditions for systems with TPMs based on temporally shuffled states. * *p* < .05, ** *p* < .01, *** *p* < .001.

The same analysis was applied to Φ calculated using the common TPM (WAKE). Similar to the results from condition-specific TPMs, Φ was highest around τ = 50 ms, with lower values observed in NREM compared to WAKE and REM (Fig. 2F). Regarding the τ yielding the maximum Φ, although ANOVA revealed a significant main effect of condition (Fig. 2G; *F(2,36)* = 183.2, *η_p_^2^* > .91, *p* < .001), post-hoc tests showed no significant differences among conditions (*ps* > .098). In contrast, the maximum Φ was significantly lower in NREM than in WAKE and REM (Fig. 2H; *F(2,36)* = 49.0, *η ^2^* > .73, *p* < .001; post hoc test, *ps* < .031). A consistent pattern of results was obtained when using length-equalized states (Fig. 2I). Furthermore, analyzing TPMs constructed from temporally-shuffled states severely diminished Φ to near-zero levels across all conditions (Fig. 2J).

### Randomized systems in V1 dataset

Fig. 3 illustrates Φ as a function of τ in randomized systems, comparing the results based on condition-specific TPMs (Fig. 3A) and a common TPM (WAKE; Fig. 3B). Across both TPM settings, Φ was highest around τ = 40–50 ms. When using condition-specific TPMs (Fig. 3A), Φ was highest in WAKE, with significant reductions observed in both NREM and REM (τ = 50 ms: *F(2,36)* = 8.5, η_p_^2^ > .32, *p* = .001, post hoc tests: *p*s < .05). In contrast, when using the common TPM (WAKE; Fig. 3B), Φ in REM recovered to levels comparable to WAKE, whereas Φ in NREM remained consistently lower (τ = 50 ms: *F(2,36)* = 31.7, η_p_^2^ > .64, *p* < .001, post hoc tests: ps < .001). These results demonstrate that state-dependent changes in Φ can be detected even in systems whose elements are constructed from random neural populations, without relying on layer-based element definitions.

**Figure 3.**
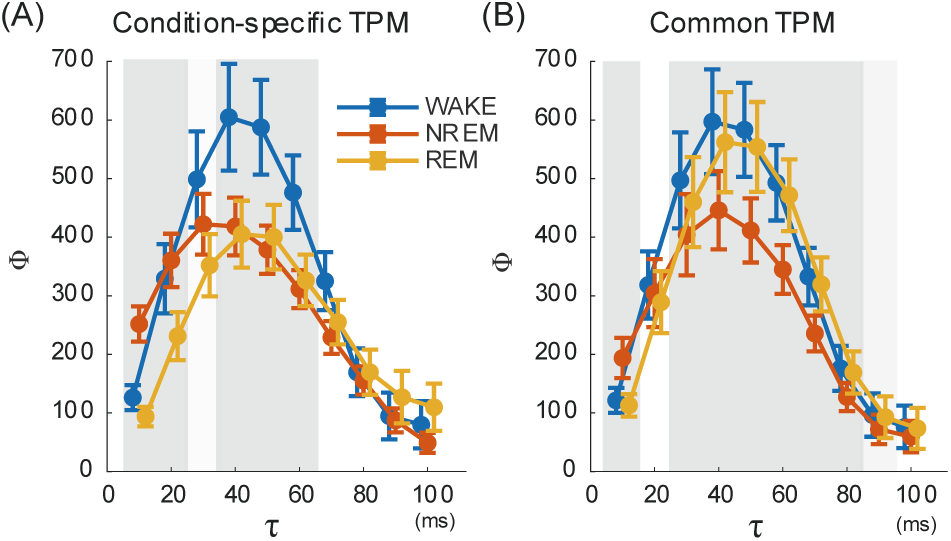
Φ as a function of τ in randomized systems for the V1 dataset. (A) Results using condition-specific transition probability matrices (TPMs). (B) Results using a common TPM (WAKE). Error bars represent the standard error of the mean across subjects. Gray shaded areas indicate τ values with a significant main effect of condition, as determined by independent ANOVAs (dark gray: Bonferroni-corrected *p* < .05, light gray: uncorrected *p* < .05).

### Effects of firing rates

Next, it was examined whether the condition-dependent changes in Φ were solely driven by neuronal firing rates. This analysis focused on Φ (τ = 50 ms) in randomized systems with a common TPM, as it exhibited the strongest relationship with firing rates. First, a comparison of the firing rate per element across conditions revealed that the firing rate in NREM was significantly lower than in WAKE and REM (Fig. 4A; *F(2,36)* = 202.97, *η_p_^2^* = .92, *p* < .001; post-hoc tests: *ps* < .001). Similarly, before controlling for firing rates, Φ was significantly lower in NREM compared to WAKE and REM (Fig. 4B; *F(2,36)* = 53.96, *η_p_^2^* = .75, *p* < .001; post-hoc tests: *ps* < .001). Correlation analysis revealed a significant positive correlation between the firing rate per element and Φ across the pooled data of all subjects and conditions (Fig. 4C; Pearson’s *r* = .36, *p* = .006), indicating that higher firing rates were associated with higher Φ. However, when analyzed separately by condition, no significant correlations were found (*r*s < .35, *p*s > .14). Furthermore, focusing on within-subject variations, the correlation between the difference in firing rates (Δ Firing Rate) and the difference in Φ (Δ Φ) was evaluated (Fig. 4D). Note that ΔΦ represents the baseline-subtracted change relative to the WAKE control; thus, negative values indicate a decrease in Φ during NREM sleep compared to WAKE. This revealed a strong positive correlation (Fig. 4D; *r* = .81, *p* < .001), demonstrating that greater reductions in firing rates correspond to greater reductions in Φ. When analyzed separately for NREM and REM, both exhibited significant positive correlations (*r*s > .63, *p*s < .003). These results suggest that the reduction of Φ in NREM could potentially be driven by the decrease in firing rates. To eliminate this confounding effect, random downsampling equalizing the firing rate per element across all conditions was applied (Fig. 4E). Recalculating and comparing Φ using this rate-controlled data revealed that, despite removing the influence of firing rates, Φ in NREM remained significantly lower than in WAKE and REM (Fig. 4F; *F(2,36)* = 36.88, *η_p_^2^* = .67, *p* < .001; post-hoc tests: *ps* < .05).

**Figure 4.**
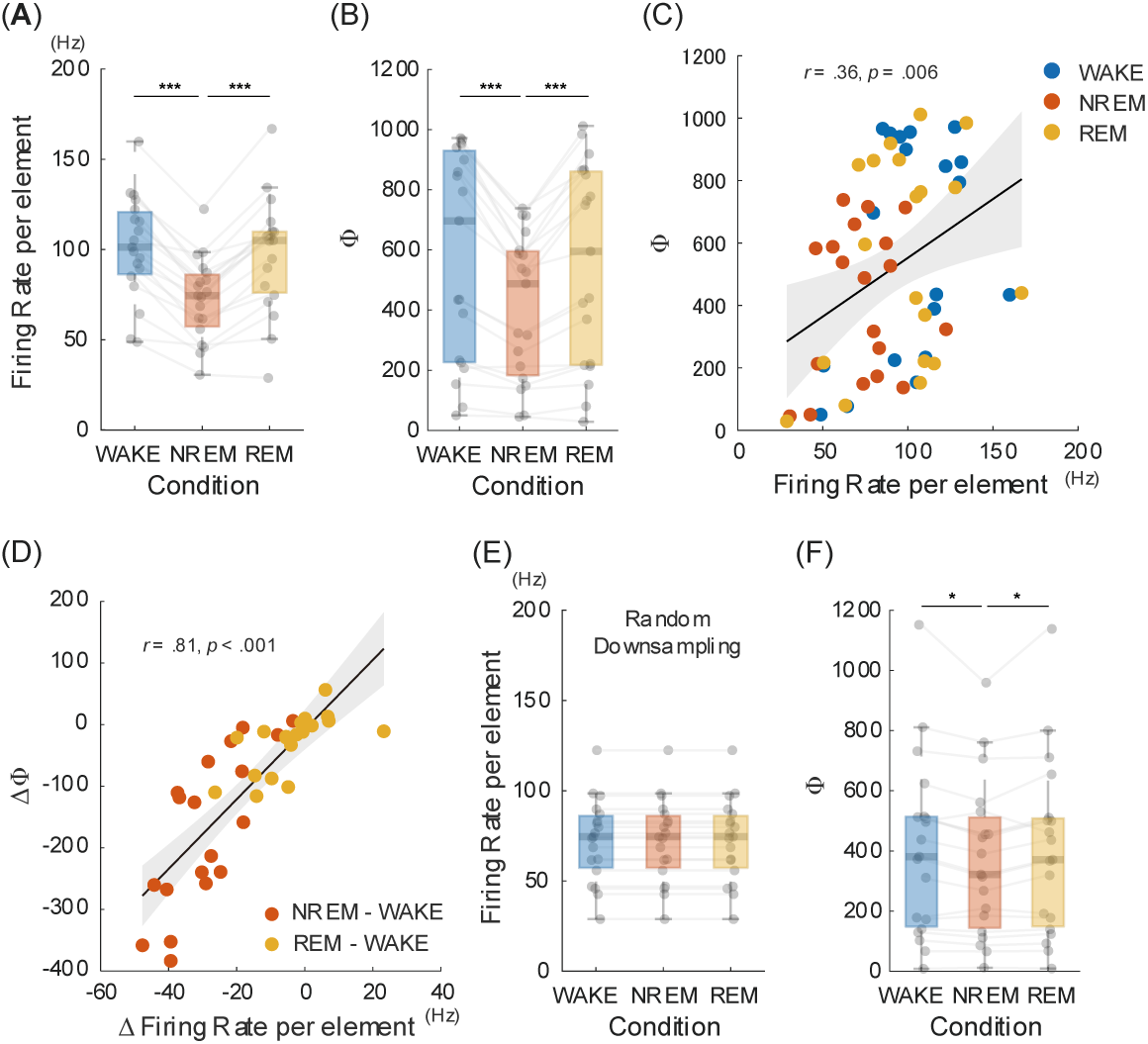
The relationship between firing rate and Φ in randomized systems and the effect of firing rate equalization. This analysis focused on Φ (τ = 50 ms) in randomized systems with a common TPM, as it exhibited the strongest correlation with the firing rate. (A) Firing rate per element across conditions. (B) Φ across conditions before firing rate equalization. (C) Scatter plot of firing rate per element and Φ across all subjects and conditions. The solid line represents the linear regression, and the shaded area indicates the 95% confidence interval. (D) Scatter plot of the within-subject differences in firing rate (Δ Firing Rate) and Φ (Δ Φ) relative to the WAKE condition (i.e., NREM - WAKE and REM - WAKE). Negative values of Δ Φ correspond to a reduction in Φ relative to WAKE. (E) Firing rate per element after applying random downsampling to equalize firing rates across conditions. (F) Φ across conditions calculated from the downsampled data. * *p* < .05, *** *p* < .001.

### Effects of ON- and OFF-periods

In NREM sleep, OFF-periods with an average duration of 175.0 ± 24.3 ms were observed intermittently. Therefore, NREM sleep was further partitioned into ON- and OFF-periods, and the mean Φ of the layer-based systems with the condition-specific TPM (τ = 50 ms) was calculated for each. Fig. 5A shows the Φ in the ON- and OFF-periods in NREM sleep. The Φ in the OFF-periods was markedly lower than that in the ON-periods (*t*(18) = 5.26, *Cohen’s d* = 1.21, *p* < .001). Fig. 5B illustrates the time course of Φ aligned to the state transitions. Upon transitioning from ON-periods to OFF-periods, Φ showed a rapid decline, whereas it exhibited a sharp increase when transitioning from the OFF-periods to the ON-periods. The duration of the Φ reduction closely coincided with the average duration of the OFF-periods.

**Figure 5.**
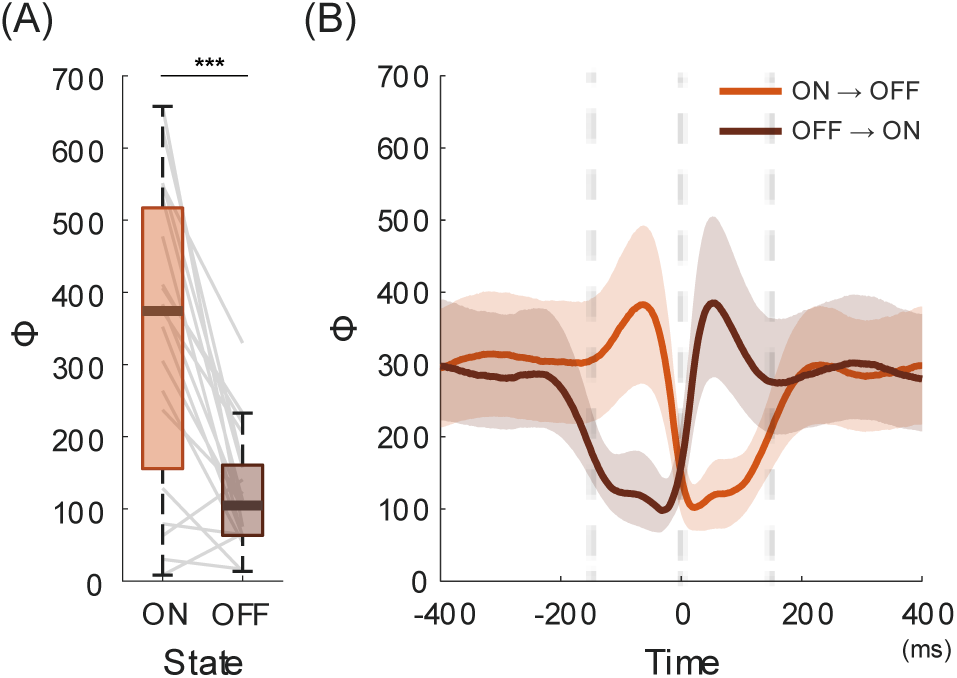
Changes in Φ across ON- and OFF-periods and peri-transition dynamics. (A) Comparison of Φ between ON- and OFF-periods. *** *p* < .001, paired t-test. (B) Time course of Φ aligned to state transition points (t = 0 ms). Vertical dashed lines at t = -175 ms and 175 ms mark the average duration of OFF-periods. The orange and brown curves indicate the transitions from ON- to OFF-periods, and from OFF- to ON-periods, respectively. Shaded areas represent the 95% confidence intervals.

### FC dataset

The analysis of the V1 dataset demonstrated that state-dependent changes in Φ can be detected in randomly clustered systems even without layer-specific structural information (Fig. 3). To test whether the local state-dependent changes in Φ observed in V1 generalize to other cortical areas, the FC dataset was analyzed (Fig. 6).

**Figure 6.**
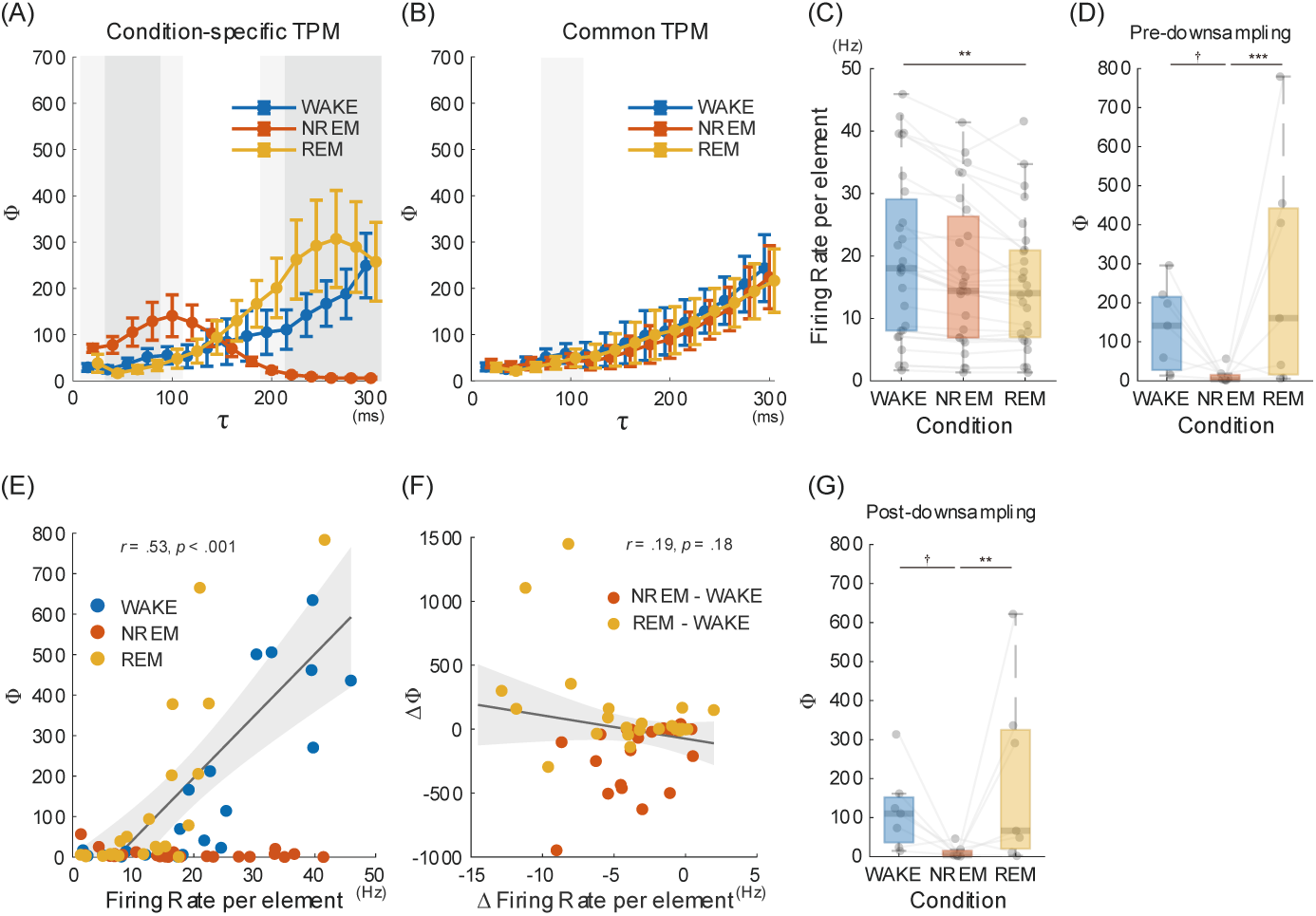
Timescale dependence of Φ, relationship with firing rates, and effects of rate-equalization via downsampling. (A) Φ as a function of the time scale τ across conditions, calculated using condition-specific Transition Probability Matrices (TPMs). Data points and error bars represent the mean ± standard error. Gray shaded areas indicate τ values with a significant main effect of condition, as determined by independent linear mixed models (dark gray: Bonferroni-corrected *p* < .05, light gray: uncorrected *p* < .05). (B) Φ as a function of τ calculated using a common TPM (WAKE). (C) Firing rate per element across conditions. Gray lines connect paired data from individual sessions. (D) Comparison of Φ across conditions prior to spike downsampling (mean value of Φ based on condition-specific TPM from 240 ms to 280 ms of τ). (E) Scatter plot of firing rate per element and Φ across all conditions. The solid line and shaded region represent the linear regression fit and 95% confidence interval, respectively. (F) Scatter plot of within-session differences in firing rate and Φ relative to WAKE. (G) Comparison of Φ across conditions after rate-equalization via random spike downsampling. † uncorrected *p* < .05, *\*\* p* < .01, *** *p* < .001.

Fig. 6A and 6B present Φ as a function of the time scale τ across conditions. Φ calculated based on condition-specific TPMs exhibited distinct profiles depending on the sleep-wake states (Fig. 6A). In the WAKE condition, Φ increased asymptotically up to τ = 300 ms. Φ in the NREM condition was highest at τ = 100 ms, whereas Φ in the REM condition reached its maximum at τ = 260 ms. At shorter time scales (τ < 100 ms), Φ during NREM was higher than that during WAKE and REM; conversely, at longer time scales (200 ms < τ < 300 ms), Φ during WAKE and REM exceeded that during NREM. In contrast, when evaluated using a common TPM (WAKE), these state-dependent differences disappeared, and Φ monotonically increased with τ across all conditions (Fig. 6B).

Next, it was examined whether these state-dependent changes in Φ depended on firing rates. Fig. 6C shows the element-wise firing rates across conditions. A significant main effect of condition was observed (*F*(2, 75) = 6.60, *η ^2^* = .15, *p* = .002), with firing rates during NREM and REM being significantly lower than during WAKE (NREM vs. WAKE: uncorrected *p* = .025; REM vs. WAKE: *p* = .002). Because the mean Φ based on condition-specific TPMs reached its maximum at τ = 260 ms in REM, Φ was averaged over the range of τ = 240–280 ms for comparison (Fig. 6D). A significant main effect of condition was identified (*F*(2, 74) = 7.72, *η ^2^* = .17, *p* < .001), with Φ during NREM being significantly lower than during WAKE and REM (NREM vs. WAKE: uncorrected *p* = .035; REM vs. WAKE: *p* < .001). Fig. 6E illustrates the scatter plot of Φ against element-wise firing rates. In data pooled across all three conditions, Φ and element-wise firing rate showed a significant positive correlation (*r* = .53, *p* < .001). Condition-wise correlation analyses revealed strong positive correlations in WAKE and REM (*r*s > .79, *p*s < .001), whereas NREM showed no significant correlation (*r* = -.29, *p* = .14). To examine the correlation between relative changes in Φ and firing rates, within-session differences relative to WAKE (NREM - WAKE, REM - WAKE) were computed. In the pooled data, no clear correlation was observed (*r* = -.19, *p* = .18; Fig. 6F). However, when analyzed separately by condition, changes in Φ and firing rates were positively correlated in NREM (*r* = .55, *p* = .003), but negatively correlated in REM (*r* = -.44, p = .023). Finally, Φ was calculated after physically controlling for firing rates using the same random downsampling procedure as applied to the V1 dataset (Fig. 6G). Even after rate-equalization, the main effect of condition remained significant (*F*(2, 74) = 5.67, *η_p_^2^* = .13, *p* = .005), and Φ during NREM remained significantly lower than during WAKE and REM (NREM vs. WAKE: uncorrected *p* = .049; REM vs. WAKE: *p* = .004).

## Discussion

This study aimed to empirically test key predictions of IIT using neural population activity from rodents. The present study revealed that integrated information Φ is highly dependent on the time window τ, highest at τ = 40–50 ms in the V1 dataset. At this optimal scale, Φ during NREM sleep was significantly lower than during WAKE. This state-dependency was robustly maintained in both layer-based and randomized systems, and persisted even after controlling for firing rates. Additionally, Φ decreased particularly during OFF-periods, showing changes that correspond to ON-OFF transitions.

Rooted in phenomenology, IIT utilizes the Φ metric to characterize the quality of conscious experience and quantify the level of consciousness in a system (Tononi et al. 2016). Previous empirical studies applying formal Φ have focused on fMRI or LFPs (Leung et al. 2021; Nemirovsky et al. 2023; Onoda et al. 2025). These studies have consistently observed reductions in global and/or local Φ during the loss of consciousness induced by sleep or anesthesia. The present study demonstrates, for the first time, that Φ similarly decreases during the loss of consciousness at the level of local neural circuits derived from spike data. Theoretical evidence suggests that the capacity of Φ to reflect state-dependent changes in consciousness is preserved even in relatively macro-level systems, where systems composed of coarse-grained elements can exhibit significant cause-effect power (Marshall et al. 2024).

One of the key findings of the present study is that Φ of the system exhibited a distinct timescale. Specifically, the observation that Φ was maximized between 40 and 50 ms in the V1 dataset aligns closely with theoretical predictions regarding the temporal grain necessary for the emergence of consciousness. According to the formulation of IIT (Albantakis et al. 2023; Marshall et al. 2018; Marshall et al. 2024), the physical substrate of consciousness is defined at the spatiotemporal resolution where Φ is maximized, as dictated by the exclusion postulate. Comolatti et al. (2025) argue that the formation of conscious experience requires a minimum temporal window (typically ranging from tens to hundreds of milliseconds) to allow for the completion of causal interactions among physical elements. These theoretical predictions are consistent with rodent neurophysiology and psychophysics. Psychophysical evidence based on critical flicker fusion thresholds indicates that the limit of the temporal integration window for visual stimuli in rodents is approximately 30 ms (Umino et al. 2018), operating on the same order of magnitude as the observed timescale. Neurophysiologically, this timescale corresponds to the optimal window for cortical coincidence detection, through which neural circuits efficiently integrate information independently of minor fluctuations in firing rates (Singer 2009). Taken together, these findings suggest that the reduction of Φ during NREM sleep is not merely a consequence of diminished neural activity but rather stems from a failure of causal integration at this optimal temporal scale. This interpretation is supported by computational simulations suggesting that Φ dynamics are not necessarily locked to firing rates (Balduzzi and Tononi 2008).

The reduction of Φ during the ON-periods can be understood from the perspective of altered network selectivity. During NREM sleep, the decrease in wake-promoting neuromodulators lowers the signal-to-noise ratio in neuronal activity (McCormick et al. 2020). Consequently, non-specific spontaneous firing increases during this period, leading to an expanded state repertoire at a microscopic level. In a coarse-grained system like the one analyzed in this study, this translates to an increase in disorganized noise, resulting in a state of high entropy. Indeed, the Shannon entropy based on the state occurrence probabilities of the 4-element system in the V1 dataset was significantly higher in the NREM condition than in both the WAKE and REM conditions (Supplementary Fig. 1). On the other hand, because the causal relationships underlying state transitions are compromised, Φ is considered to be reduced during NREM sleep.

Furthermore, the bistable dynamics (ON and OFF periods) during NREM sleep play a critical role in the fragmentation of consciousness. In the present study, while Φ was reduced even during ON-periods compared to the WAKE condition, the reduction during OFF-periods was particularly evident. This twofold reduction is consistent with computer simulations demonstrating that systems governed by bistable dynamics cannot sustain high Φ, as it collapses near zero both during highly active states (ON-periods) and inactive states (OFF-periods) (Balduzzi and Tononi 2008). The decline in Φ arises because cause-effect chains are physically interrupted by the forced insertion of OFF-periods, which likely shares the same physiological mechanism as the reduction in perturbational complexity observed following transcranial magnetic stimulation during NREM sleep (Massimini et al. 2005). At the physiological level, the diminished release of arousal-promoting neuromodulators during NREM sleep alters membrane conductance, predisposing neurons to hyperpolarization and rendering them unable to sustain firing for durations exceeding several hundred milliseconds (Steriade et al. 2001). Information-theoretically, these population-wide OFF-periods collapse the system into a silent state, thereby severely restricting the repertoire of states and diminishing the cause-effect power required for high Φ.

Another finding of the current study is that local circuit Φ remained high during REM sleep comparable to that during WAKE. This result aligns well with the prediction of IIT, which posits that Φ tracks the presence of conscious experience rather than mere behavioral responsiveness or external sensory input. REM sleep is accompanied by vivid, internally generated conscious experiences (Nir and Tononi 2010). The elevated Φ during REM sleep suggests that internal causal integration within local cortical circuits is preserved during dream states. The preservation of high Φ during REM sleep can be attributed to the restoration of cortical activation and the absence of OFF-periods. Unlike NREM sleep, REM sleep is characterized by a desynchronized, wake-like electroencephalographic pattern driven by high cholinergic activity (Steriade et al. 2001), which prevents local populations from falling into OFF-periods. Consequently, the network avoids the loss of state-transition dynamics observed during NREM sleep, thereby maintaining the cause-effect power required for high Φ.

The observed reduction of Φ during NREM sleep in both V1 and FC suggests that the loss of consciousness involves a widespread transformation of causal structures at the local circuit level. Interestingly, a previous fMRI study (Onoda et al. 2025) reported that while Φ decreased specifically in the posterior cortex during NREM sleep, no significant changes were found in prefrontal or whole-brain networks. This discrepancy may arise because the highly diverse and sparse nature of FC neural activity (Le Merre et al. 2026) complicates the detection of integrated information at the macro-imaging scale. In higher-order regions, the fine-grained causal structures that define Φ may be averaged out within coarse fMRI voxels, leading to an underestimation of its reduction during sleep. Furthermore, the longer timescale observed in the FC compared to V1 likely reflects the functional hierarchy of the cortex (Gao et al. 2022). As proposed by Marshall et al. (2024), the macro spatiotemporal grain at which a system maximizes its causal power is not fixed but varies across regions. While V1 relies on rapid neural dynamics suited for early sensory processing, higher-order regions like the FC require longer intrinsic timescales to integrate and maintain information. From the perspective of IIT, the longer timescale in the FC suggests that this region integrates causal interactions over a broader temporal window, supporting the complex structures necessary for higher-order awareness and reportability (Koch et al. 2016; Odegaard et al. 2017; Seth and Bayne 2022).

A notable discrepancy observed in this study is the behavior of Φ in the FC dataset under the common TPM versus condition-specific TPMs (Fig. 6). While V1 exhibited a consistent collapse of Φ during NREM sleep regardless of TPM specification, the reduction of Φ in FC was significantly detected only when condition-specific TPMs were used. This discrepancy suggests the regional specificity of cortical causal structures during sleep. As previously noted, dense neuromodulatory inputs undergo suppression during NREM sleep (McCormick et al. 2020), a feature that is pronounced in the FC (Brown et al. 2012). Such neurochemical shifts can fundamentally alter synaptic efficacy, implying that the underlying causal mechanism (i.e., TPM) reconfigures between WAKE and NREM in FC. Consequently, applying a WAKE-derived common TPM to NREM state sequences fails to capture NREM-specific causal constraints, thereby obscuring the state-dependent difference in Φ. In addition, unlike V1, which is strongly constrained by stereotyped local feedforward architecture (Douglas and Martin 2004), FC circuits possess higher metastability and complex recurrent dynamics (Murray et al. 2014). In such high-order networks, the loss of integrated information during NREM sleep may reflect a breakdown of state-transition dynamics embedded in the state-dependent TPM rather than simple shifts in state occupation probability.

Several limitations of the present study should be acknowledged. First, the analysis was restricted to systems composed of only four elements. While this constraint was necessary due to the exponential increase in computational complexity associated with Φ calculation, it precluded the systematic search for the physical substrate of consciousness within larger neural populations. Consequently, the observed changes in Φ may represent only a partial reflection of the causal dynamics occurring within the broader cortical network. Second, the findings in this study are based on recordings from specific cortical regions, namely the mouse V1 and rat FC. While these areas are central to sensory processing and higher-order cognition, the loss of consciousness is a global phenomenon involving complex interactions between the cortex and subcortical structures, such as the thalamus. Because the present study did not account for these thalamocortical loops, the observed reduction in Φ should be interpreted as a local manifestation of a more extensive, system-wide disruption of causal integration.

In conclusion, the present study provides empirical support consistent with predictions of IIT using neural population activity from the mouse V1 and rat FC. The primary finding was the reduction of Φ during NREM sleep at the local circuit level. This phenomenon was consistently observed not only in layer-based systems but also in randomized systems, suggesting that the loss of consciousness involves a widespread transformation of causal structures that are independent of specific circuit architectures. Furthermore, the finding that Φ is maximized at a specific time window τ supports the theoretical requirement of a temporal grain necessary for the emergence of consciousness. The tendency for the timescale to be longer in the FC likely reflects distinct intrinsic timescales associated with the cortical hierarchy. Future research exploring causal structures in larger-scale networks will be essential to elucidate the physical substrate of consciousness.

## Funding

This work was supported by JSPS KAKENHI Grant Number 23K03022.

## Conflict of interest statement

The author declares no conflicts of interest.

## Acknowledgments

This work was supported by JSPS KAKENHI Grant Number 23K03022.

---

**Supplementary Figure 1.**
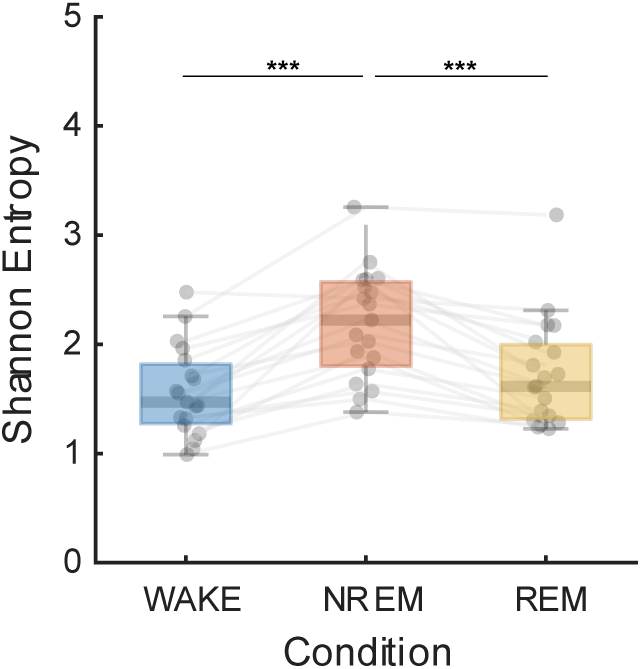
Shannon entropy of system states across conditions for the V1 dataset.

## Notes

### Competing Interest Statement

The authors have declared no competing interest.

### Summary of Updates

Introduction revised to explicitly define micro- and macro-scales and clarify the rationale for macro-units based on IIT 4.0 Intrinsic Units; additional control analyses performed for TPM estimation (temporal shuffling, state-matching, and condition-specific TPMs); spike downsampling analysis added to equalize firing rates; timescale (τ) search range extended up to 300 ms for the FC dataset; normalized Φ replaced with raw Φ throughout; Discussion updated regarding ON/OFF period dynamics and Shannon entropy; Supplementary Figure 1 added; overstatements and terminology (stable Φ, peak Φ) refined throughout.

